# Nine Novel Cluster E Mycobacteriophages Isolated from Soil

**DOI:** 10.1101/2020.03.05.979757

**Authors:** Joseph M. Gaballa, Amanda Freise, Jordan Moberg Parker

## Abstract

Novel mycobacteriophages FireRed, MISSy, MPhalcon, Murica, Sassay, Terminus, Willez, YassJohnny, and Youngblood were isolated from soil samples on the host *Mycobacterium smegmatis*. Transmission electron microscopy revealed phage structures suggestive of the *Siphoviridae* family of phages. Phage genomes were sequenced and annotated, revealing that they belong to actinobacteriophage Cluster E. Here, we describe the features of these phage genomes and discuss their similarity to previously discovered cluster E phages.

## INTRODUCTION

Mycobacteriophages represent a diverse class of viruses that selectively infect bacterial hosts from the genus *mycobacterium*. The past decade has seen a rapid increase in the discovery and genomic annotation of mycobacteriophages, many of which are of translational interest because of their ability to infect and lyse the bacterial pathogens *Mycobacterium tuberculosis* and *Mycobacterium leprae* (1). The discovery of novel mycobacteriophages is typically conducted using *Mycobacterium smegmatis*, a non-pathogenic strain that serves a model host for evaluating a mycobacteriophage’s therapeutic potential. Over 150,000 genes have been identified in the collection of over 1,800 mycobacteriophages which have been sequenced (2). This incredible degree of genetic diversity warrants mycobacteriophages to be organized into clusters, and further subclusters, based on genomic similarity (3). Our group’s previous work focused on structurally and genomically characterizing novel Cluster K mycobacteriophages isolated from environmental soil samples (4). In the present study, we introduce nine novel Cluster E mycobacteriophages isolated from soil samples. Transmission electron microscopy (TEM) revealed siphoviral morphology, while genome sequencing and annotation indicated a high degree of similarity between the genomes of the novel phages and previously isolated Cluster E phages.

## METHODS

### Phage isolation

Soil samples were collected from sites around the greater Los Angeles, CA area. Phages were isolated using direct or enrichment isolation on *Mycobacterium smegmatis* mc2155 as described by the SEA-PHAGES Discovery Guide (https://seaphagesphagediscoveryguide.helpdocsonline.com/). Briefly, for direct isolation, soil samples were suspended in 1X 7H9 broth supplemented with albumin dextrose and incubated on shaker at room temperature for 1-2 hours, followed by filtration to remove bacteria. For enrichment isolation, sterile water was added to soil in a flask along with 10X 7H9 broth supplemented with albumin dextrose and *M. smegmatis* mc2155 culture. The enrichment culture was incubated on a shaker for 24 hours at room temperature filtered to remove bacteria. Filtrates were spotted onto lawns of *M. smegmatis* and incubated at 35°C to check for clearing. Successive plaque assays were performed on filtrates with clearings to isolate and amplify phages for further characterization.

### Transmission electron microscopy

Purified phage lysates were stained with 1% uranyl acetate on carbon-coated grids and visualized with a transmission electron microscope at magnifications ranging from 21,000-67,000X.

### Genome sequencing, assembly, and annotation

Viral DNA was isolated using the Wizard Promega DNA Clean-Up Kit (Product #A7280) and sequenced via Illumina or 454 pyrosequencing (Table 1). Location and coding potential of putative genes were predicted using DNA Master (J. G. Lawrence lab [http://cobamide2.bio.pitt.edu]), which integrates both Glimmer and GeneMark to detect potential open-reading frames (5, 6). Location calls were curated using Phamerator, which sorts genes into “phamily” groups based on pairwise amino acid comparisons, and Starterator, which identifies conserved start sites (7). ARAGORN was used to detect the presence of tRNA genes (8). Putative functional calls were predicted using BLASTp against PhagesDB and NCBI databases (9). NCBI’s Conserved Domain Database, HHPred, and TMHMM were also used to assign putative functions (10–12).

**Table 1:** Novel Cluster E Phage Isolation and Sequencing Data.

| **Mycobacteriophage** | **Isolation method** | **Year of Isolation** | **Sequencing Site** | **Sequencing Coverage** |
| --- | --- | --- | --- | --- |
| FireRed | Enriched | 2013 | The UCLA Genotyping and Sequencing Core | 48x |
| MISSy | Enriched | 2014 | The UCLA Genotyping and Sequencing Core | 63x |
| MPhalcon | Direct | 2017 | Pittsburgh Bacteriophage Institute | 2029x |
| Murica | Enriched | 2013 | The UCLA Genotyping and Sequencing Core | 81x |
| Sassay | Enriched | 2014 | The UCLA Genotyping and Sequencing Core | 126x |
| Terminus | Enriched | 2014 | Illumina Sequencing | 10,582x |
| Willez | Enriched | 2011 | The UCLA Genotyping and Sequencing Core | 157x |
| YassJohnny | Enriched | 2015 | Pittsburgh Bacteriophage Institute | 913x |
| Youngblood | Enriched | 2014 | The UCLA Genotyping and Sequencing Core | 38x |

## RESULTS

### Phage Phenotypes

Purification and amplification of the phages revealed heterogenous plaque morphologies, including clear, turbid, “halo”, and “bullseye” morphologies with average diameters ranging from 2mm-6mm (Figure 1). Transmission electron microscopy revealed that all of the phages had flexible non-contractile tails and icosahedral capsids (Figure 2).

**Figure 1:**
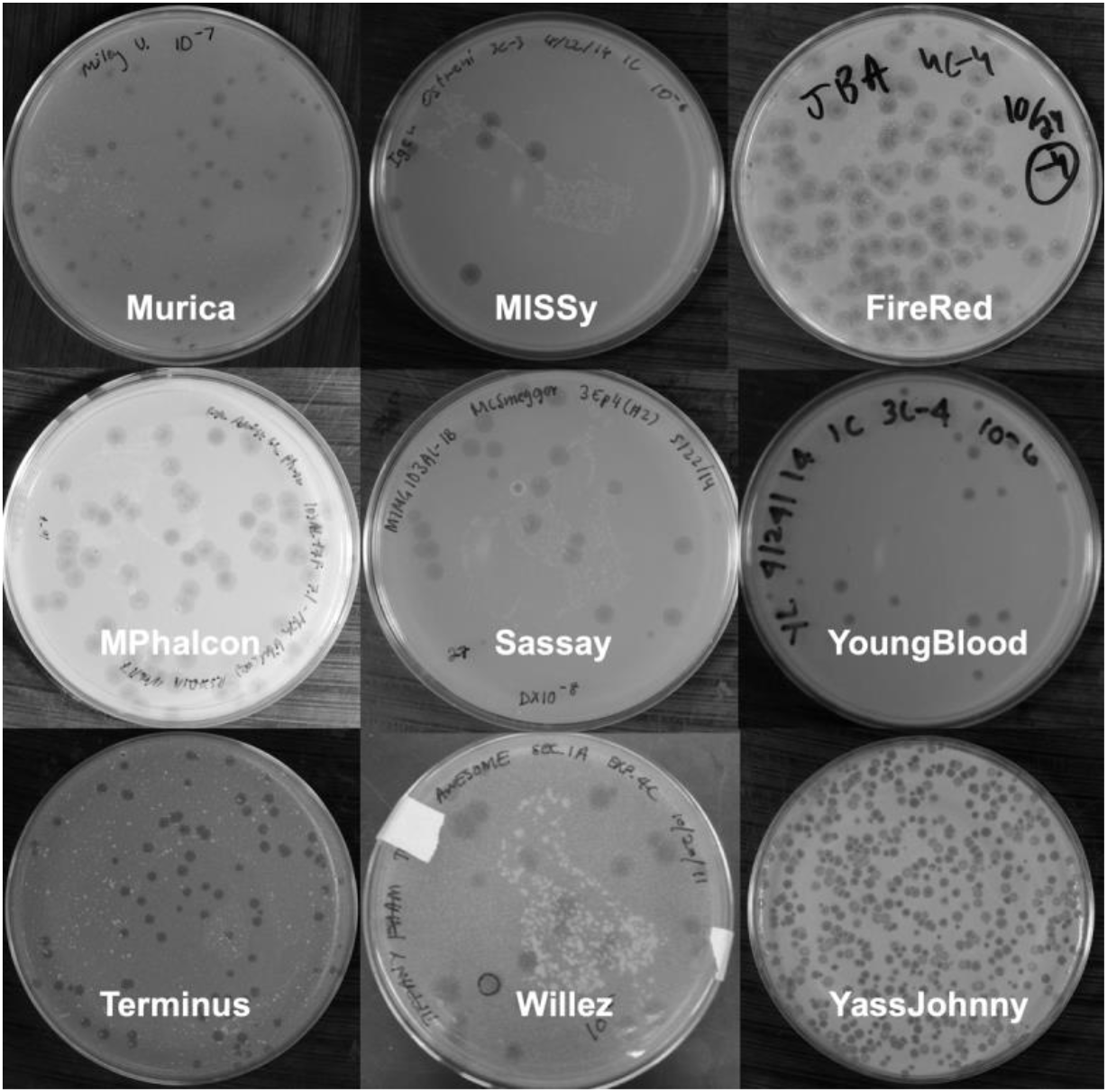
Plaque morphologies of Cluster E phages. Representative 90mm agar plates after successive rounds of viral propagation in *M. smegmatis*.

**Figure 2:**
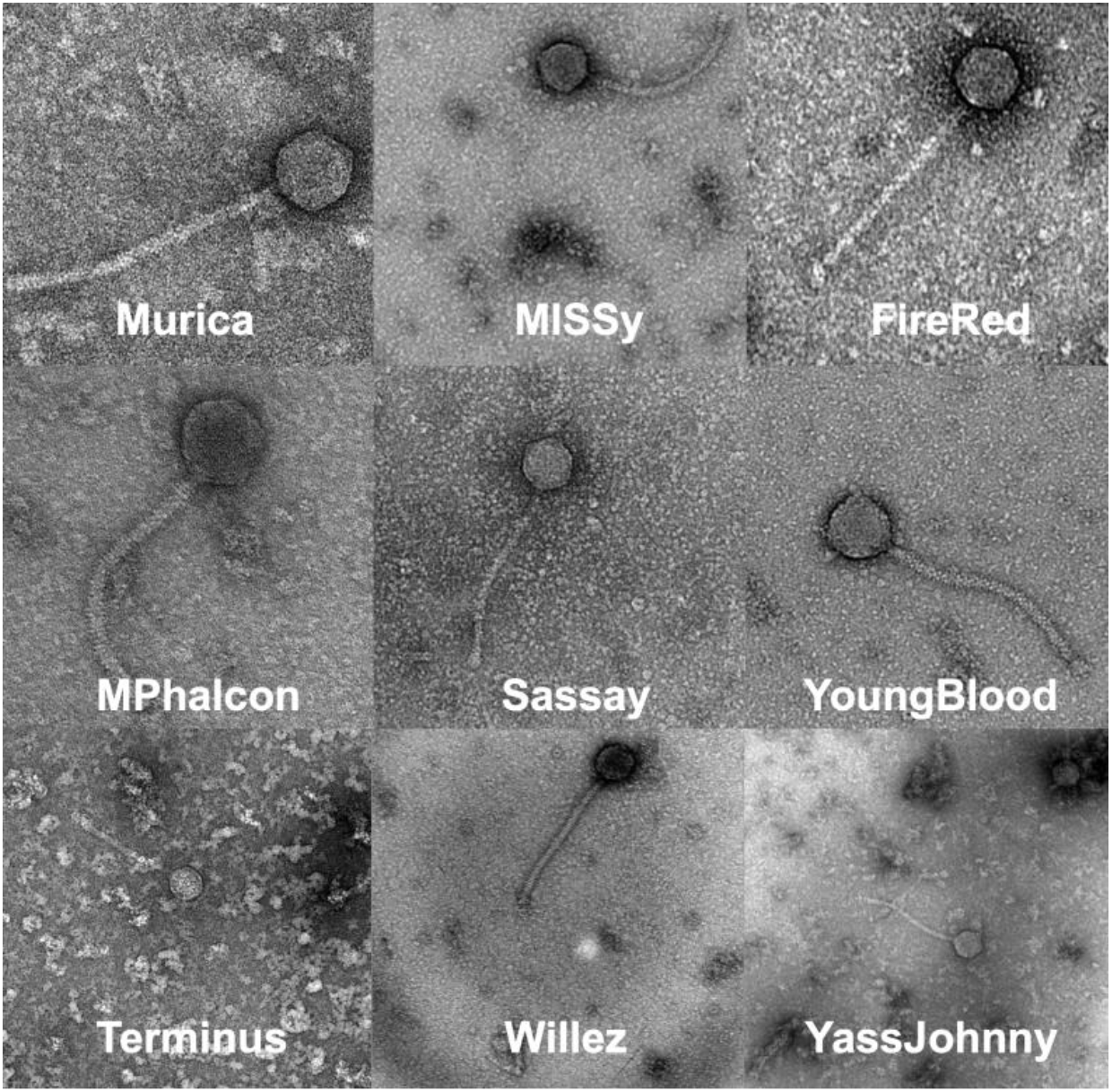
Transmission electron microscopy of novel Cluster E phages. Representative EM pictures taken at magnifications ranging between 21,000-67,000X. Key features of viral morphology include flexible, non-contractile tails and icosahedral heads indicative of *Siphoviridae* family members.

### Genomic Analysis

Genome lengths ranged between 73,495 and 77,053 base pairs with each genome containing a 3’ sticky overhang (CGCTTGTCA). GC content ranged from 62.9% to 63.1%. Total gene count ranged between 141-150 predicted genes (Table 2). A Phamerator map was generated to compare the genomic architecture of the nine phages, which indicated that regions containing approximately 15-18 reverse-coding genes at the tail end of the genome are consistent among the phages (Figure 3). Each phage genome contains two tRNAs, and all of the genomes contain a lysis cassette comprising Lysin A, Lysin B, and holin genes. Integrase and an immunity repressor are found downstream of the lysis cassettes in all phages.

**Table 2:**
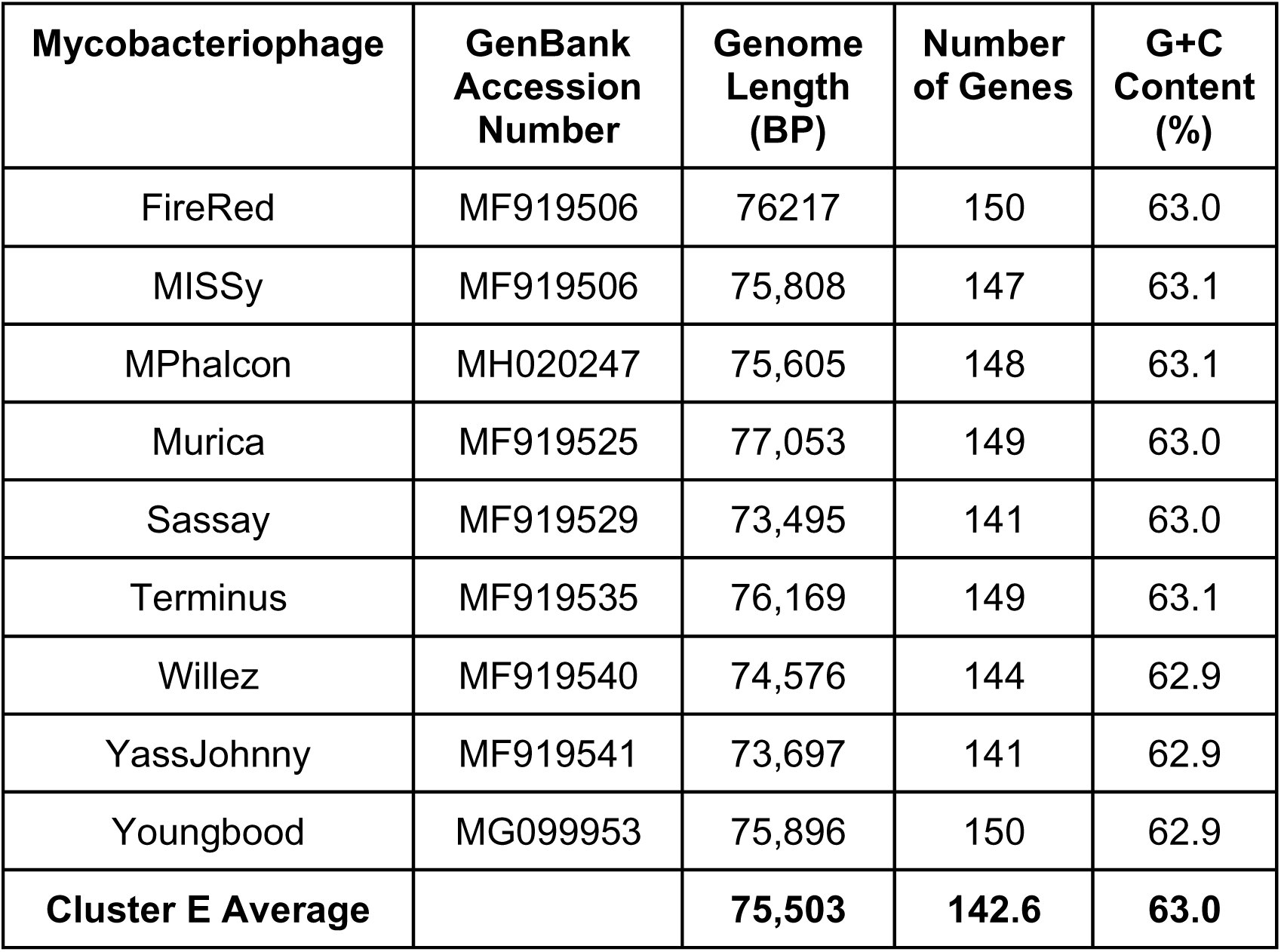
Novel Cluster E Phage Genomic Data.

**Figure 3:**
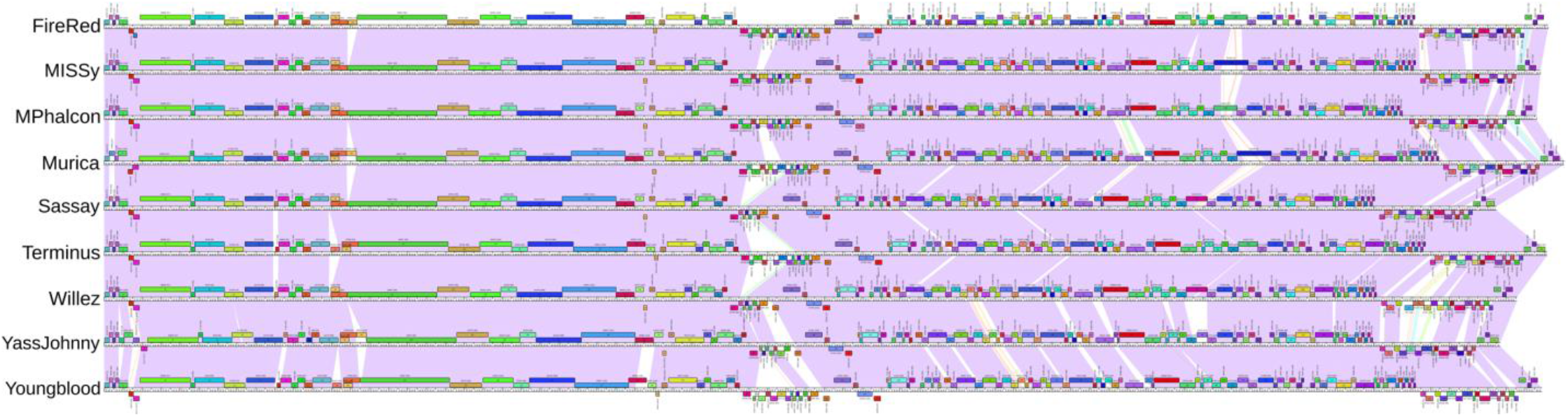
Phamerator map of Novel Cluster E phages. Comparative genomic analysis reveals similar genomic architecture among the discovered phages. The most variability between phages and the highest concentration of genes with no known functions are found on the right sides of the genomes.

## DISCUSSION

Nine bacteriophages capable of infecting *M. smegmatis* were discovered in soil samples. Upon examining the structure of the phages with TEM, features consistent with the *Siphoviridae* family of phages such as long non-contractile tails and icosahedral heads were seen. Based on nucleotide similarity, these phages belong to Cluster E, which had 102 members at time of publication (phagesdb.org). The average GC-content of these phages (63.0%) is consistent with the average of published Cluster E phages (63.0%).

The novel phages contain between 141 and 150 predicted genes, with seven out of nine phages expressing more than the average gene count for Cluster E phages (142.6). As is common in Cluster E phages, most of the genes with no known functions are located on the right side of the genome, while the left side of the genome is highly conserved and encodes most of the structural proteins.

The presence of a lysis cassette, integrase, and an immunity repressor suggests that these phages can undergo lytic or lysogenic lifecycles, and this is supported by the presence of both clear and turbid or bullseye plaque morphologies that the most phages display. Nonetheless, additional experiments need to be done to confirm the ability of each phage to form lysogens.

## Acknowledgements

Mycobacteriophages were isolated by undergraduates in the Research Immersion in Virology course-based undergraduate research experience in the Microbiology, Immunology, and Molecular Genetics Department at UCLA. This project was funded by the Dean of Life Sciences Division at UCLA, with additional support for sequencing from the HHMI Science Education Alliance-Phage Hunters Advancing Genomics and Evolutionary Science (SEA-PHAGES) program. We thank Krisanavane Reddi for preparation of materials, lysate archiving, and management of the instructional laboratories; Hong Zhou of the UCLA Electron Imaging Center for NanoMachines for electron microscopy support; Rebecca A. Garlena and Daniel A. Russell at the Pittsburgh Bacteriophage Institute for phage sequencing and assistance with genome assembly; and Debbie Jacobs-Sera, Welkin Pope, Graham Hatfull, and the SEA-PHAGES community for programmatic support.

## References

1. Gondil VS., Chhibber S. 2018. Exploring potential of phage therapy for tuberculosis using model organism. Biomed Biotechnol Res J, 2(1), 9–15.

2. Hatfull G. 2018. Mycobacteriophages. Microbiol Spectrum 6(5):GPP3–0026-2018. doi:10.1128/microbiolspec.GPP3-0026-2018.

3. Hatfull, G.F., Jacobs-Sera, D., Lawrence, J.G., Pope, W.H., Russell, D.A., Ko, C., Weber, R.J., Patel, M.C., Germane, K.L., Edgar, R.H., Hoyte, N.N., Bowman, C.A., Tantoco, A.T., Paladin, E.C., Myers, M.S., Smith, A.L., Grace, M.S., Pham, T.T., O'Brien, M.B., Vogelsberger, A.M., Hryckowian, A.J., Wynalek, J.L., Donis-Keller, H., Bogel, M., Peebles, C.L., Cresawn, S.G., & Hendrix, R.W. 2010. Comparative genomic analysis of 60 Mycobacteriophage genomes: genome clustering, gene acquisition, and gene size. Journal of molecular biology, 397(1), 119–43.

4. Genome Sequences of Cluster K Mycobacteriophages Deby, LaterM, LilPharaoh, Paola, SgtBeansprout, and Sulley. Gaballa JM, Dabrian K, Desai R, Ngo R, Park D, Sakaji E, Sun Y, Tan B, Brinck M, Brobst O, Fernando R, Kim H, McCarthy S, Murphy M, Sarkis A, Sevier P, Singh A, Wu D, Wu MY, Ennis HA, Luhar R, Miller JE, Orchanian SB, Salbato AN, Alam S, Brenner L, Kailani Z, Laskow J, Ma X, Miikeda A, Nol-Bernardino P, Sukhina A, Walas N, Wei W, Do NP, Fournier CT, Kim CJ, Mosier SF, Pierson C, Romero IG, Sanchez M, Sawyerr O, Wang J, Watanabe R, Wu S, Chen A, Kazane K, Kettoola Y, Goodwin EC, Lund AJ, Villella W, Williams D, Freise A, Moberg Parker J. Microbiology Resource Announcements. 2019. PMID: 30643892

5. Salzberg, S. L., Delcher, A. L., Kasif, S., & White, O. 1998. Microbial gene identification using interpolated Markov models. Nucleic Acids Research, 26(2), 544–548.

6. Lukashin, A. V., & Borodovsky, M. 1998. GeneMark.hmm: new solutions for gene finding. Nucleic Acids Research, 26(4), 1107–1115.

7. Cresawn SG, Bogel M, Day N, Jacobs-Sera D, Hendrix RW, Hatfull GF. 2011. Phamerator: a bioinformatic tool for comparative bacteriophage genomics. BMC Bioinformatics, 12(395). 10.1186/1471-2105-12-395.

8. Laslett, D., & Canback, B. 2004. ARAGORN, a program to detect tRNA genes and tmRNA genes in nucleotide sequences. Nucleic Acids Research, 32(1), 11–16.

9. Altschul, S.F., Gish, W., Miller, W., Myers, E.W. & Lipman, D.J. 1990. “Basic local alignment search tool.” J. Mol. Biol., 215, 403–410.

10. Marchler-Bauer, A., & Bryant, S. H. 2004. CD-Search: protein domain annotations on the fly. Nucleic Acids Research, 32(Web Server issue), W327–W331. http://doi.org/10.1093/nar/gkh454.

11. Johannes Söding, Andreas Biegert, Andrei N. Lupas. 2005. The HHpred interactive server for protein homology detection and structure prediction. Nucleic Acids Research, 33(2), 244–248. https://doi.org/10.1093/nar/gki408

12. Krogh A, Larsson B, von Heijne G, Sonnhammer ELL. 2001. Predicting transmembrane protein topology with a hidden Markov model: Application to complete genomes. Journal of Molecular Biology. 305(3), 567–580.

